# A novel agar-based millifluidic device for spatially-structured microbial populations

**DOI:** 10.64898/2026.06.11.731666

**Authors:** Brandon Tuck, Stefano Pagliara, Wolfram Möbius

**Affiliations:** Living Systems Institute, Faculty of Health and Life Sciences, University of Exeter, Exeter, UK; Biosciences, Faculty of Health and Life Sciences, University of Exeter, Exeter, UK; Physics & Astronomy, Faculty of Environment, Science and Economy, University of Exeter, Exeter, UK

## Abstract

The role of spatial structure in microbial ecology and evolution is increasingly recognised and investigated, often with agar plates as a template for a spatially structured environment. While convenient, agar plates do not allow for the spatial and temporal control microbiologists have become accustomed to in the field of microfluidics with its tight environmental control for single cells and small populations, holding back research on surfaces and at larger length scales and population sizes.

To close this gap, we developed a novel device with an agar sheet sealing indented channels through which media perfuses. As proof of principle, we grew populations of non-motile *Escherichia coli* and motile *Pseudomonas aeruginosa* for 60 hours with continuous propagation of the colony’s front, in contrast to agar plates where growth declined much earlier and stopped after about 40 hours. To demonstrate the capabilities of spatial control, we grew *P. aeruginosa* along different temporally-stable gradients of the cephalosporin antibiotic ceftazidime and characterised the emerging bacterial growth patterns.

The device is a step towards highly controlled studies of microbial populations in continuous, non-uniform spatially structured environments. Designed with cost and accessibility in mind, we believe that this novel device will enable new insights into microbial ecology and evolution.

## Introduction

Culturing bacteria on the surface of an agar sheet with growth medium is a well-established laboratory technique and the primary means to establish clonal populations in microbiology [1]. Growth on an agar surface arguably also serves as the most widely used assay to investigate the role of spatial structure in microbial ecology and evolution, for example in the study of biofilms [2]. Ease of use and affordability of agar sheets in Petri dishes are key advantages, but the inability to control environments spatially and temporally is a major disadvantage.

Varying medium composition and concentration of agar are key experimental variables affecting the growth of microbial populations [3], but the environment cells experience is not stable unless cells are transferred to a new surface, for example by replica plating [4]. Keeping the environment stable by increasing the size of the agar sheet and providing a reservoir is not a feasible strategy because diffusion is too slow on the scale of the agar sheet to provide a constant environment. The characteristic diffusion time for a glucose molecule with diffusion coefficient of 6 *·* 10^−6^ cm^2^/s [5] diffusing through water and across 1 mm of a low concentration agar slab is under an hour and across 1 cm is under a day. However, it would take weeks to diffuse from the boundary to the centre of a standard agar sheet. Nutrient depletion due to bacterial uptake and accumulation of potential toxic byproducts produced via cell metabolism are therefore typically localised to the site of growth and contribute to an increasingly unfavourable local environment over time [6, 7]. For populations of non-motile cells, this eventually halts growth altogether, limiting colony size.

While diffusion is too slow for a colony to benefit from nutrients across the whole agar plate, it is too fast to maintain spatial patterns at the scale of typical populations on the surface. Consider a gradient of antibiotics over 1 cm; it will change substantially while the population grows and expands over the course of hours and days. This shortcoming has been addressed by dramatically increasing system size and using a model system of chemotaxing bacteria [8]; however, this approach is not transferable to other systems and requires a lot of resources. Taken together, diffusion is limiting experiments as it is too slow to provide a truly constant environment and too fast to maintain spatial variation at the length and time scales of interest.

Short of transferring populations to new environments such as replica plating, a sheet of agar in a Petri dish does not allow for temporal control. This is different for small populations or even single cells. Microfluidic platforms [9–16] allow for spatial as well as fast temporal control over the microbial environment. Cells are confined to a single layer to allow visualisation of single cells moving across a plane [17, 18] or multiple layers to resemble biofilms [19–22]. Spatial structuring allows one to study bacterial chemotaxis [23–25] or the evolution in gradients of antibiotics [26].

The experimental opportunities microfluidic platforms offer and the deficiency of methods to study microbial populations on surfaces at the scale of millimeters to centimeters encouraged us to develop a device that brings concepts from microfluidics to the classic agar sheet to allow for new quantitative research in microbial ecology and evolution. We thereby aimed for simplicity and affordability inspired by recent work on affordable process control techniques for macroscopic well-mixed microbial populations [27, 28] and used finite-element simulations to guide the design. Following an outline of design considerations and device fabrication, we provide two proof of concept applications, population growth in a spatially uniform environment and growth on an antibiotic gradient.

## Materials and Methods

### Strains and media

Strains used in this study were *E. coli* strain DH5*α, P. aeruginosa* strain PA14 (labelled *P. aeruginosa* PS (i.e., parental strain) in this publication) as well as *P. aeruginosa* strain PA14 flgK::Tn5B30(TcR) [29] (labelled *P. aeruginosa* Δ*flgK* here). *P. aeruginosa* strains are the same to those used by Dimitriu *et al*. [30].

LB growth medium (10 g/L tryptone, 5 g/L yeast extract and 10 g/L NaCl) was used for overnight culture and flow within the device. LB agar plates (LB with 1.5% w/v agar) were used for clonal culture. Where ceftazidime (Sigma Aldrich) was used in the device, it was added in powder form to LB growth medium to the desired concentration.

### Device components

The primary device assembly consists of three main components. The core is a polymer support with embedded channels, connected via channel inlets and outlets to plastic tubing, where the inlet tubes are connected to reservoir flasks and the outlet tubes to waste syringes. This assembly is then combined with the agar sheet to perform experiments (Fig. 1).

**Figure 1:**
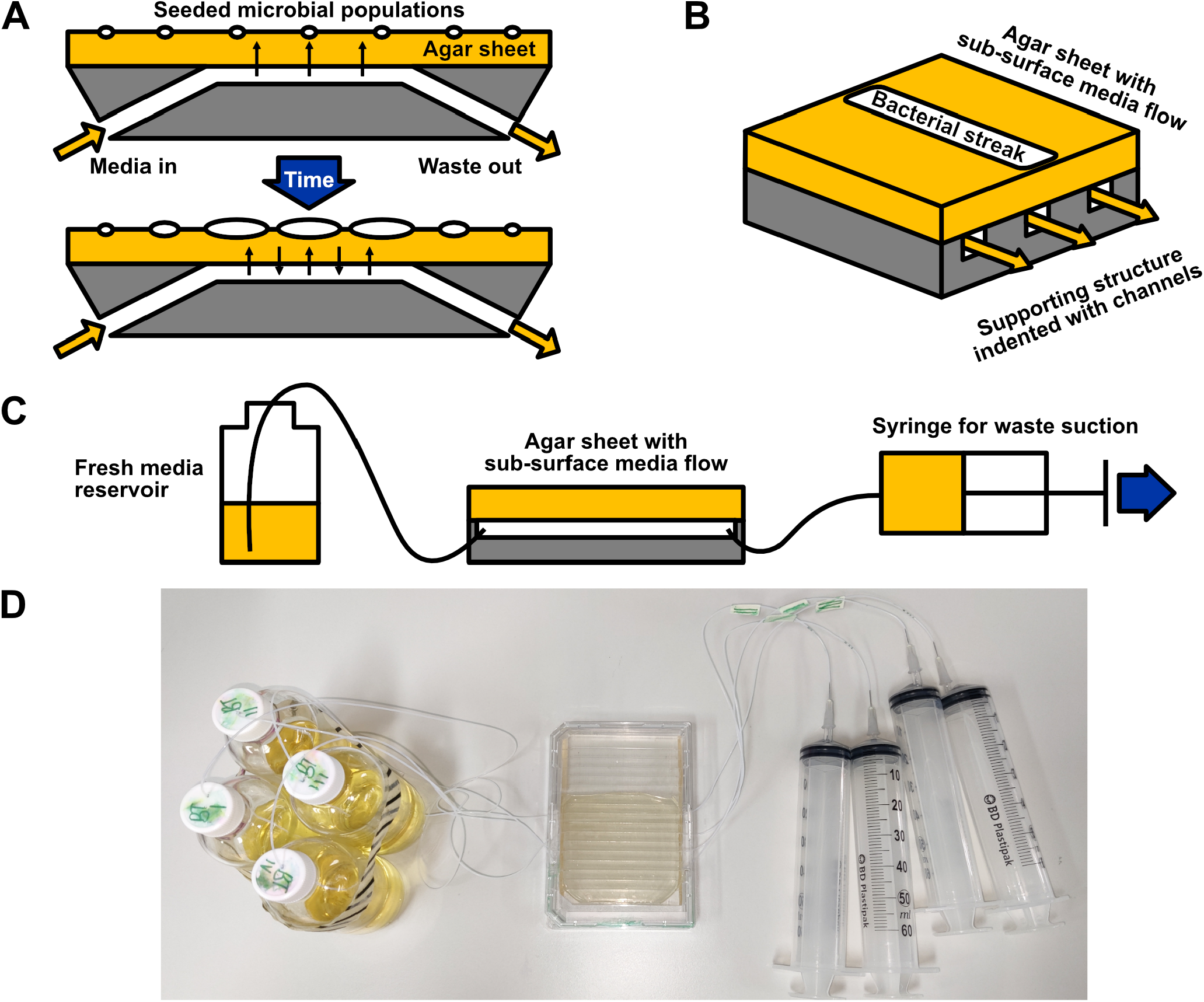
Overview of the agar-based fluidic device and its function. (A) Controlled flow of growth media under an agar surface allows molecules, e.g., nutrients, to diffuse vertically through the agar towards the surface. Bacteria seeded directly above the channel have constant access to nutrients over time. Conversely, metabolic byproducts diffuse into the fluid stream and are removed from the microbial environment. Over longer time frames, areas outside the direct vicinity of the channel also become viable for growth as nutrients diffuse from adjacent channels allowing sustained population expansion. (B) An isometric view of the device layout with a streak of bacteria on the surface. Channels used are 1 mm in width and height and spaced 5 mm apart. Agar thickness is 1.7 mm. (C) Set-up of the device with input and output reservoirs: a bottle of media supplies the device and a syringe collects waste, kept under negative pressure using a pump for constant suction. The total channel length is 70 mm. D) Photo of the set-up from an aerial view.

The polymer support is made from polydimethylsiloxane (PDMS), a transparent and autoclavable elastomer and the most common material for devices in microfluidics for biological applications. Due to the larger scale of the device, the template for the support can be 3D-printed rather than etched into silicon, reducing need for specialised equipment. We used a stereolithography 3D printer for high resolution with printing resin capable of supporting the required 70 °C PDMS curing process (Formlabs Form 3 printer with Formlabs standard clear resin). The tubing used was made of polytetrafluoroethane (PTFE) (see Ref. [31] for more details), an autoclavable polymer. Tubing were connected to reservoir bottles made of borosilicate glass.

Thin agar sheets were prepared by casting nutrient-free agar (1.5% w/v agar) in a sterile 14 cm borosilicate glass Petri dish (Fisher Scientific). We found agar sheets that were poured and refrigerated within hours to lack sufficient adhesion with PDMS to produce a functional device, as leakage of perfused media was a common phenomenon. This issue could be remedied by conditioning the agar sheet prior to device assembly. The conditioning step involved drying the agar sheets at 37 °C for 8 to 24 hours depending on the ventilation in the dish. This step increased adhesion, arguably due to changing the surface properties of the agar.

### Device assembly and growth of microbial populations on the device

We prepared cultures by picking a single colony from a streak plate and transferring it into growth tubes with 5 mL of LB media in a 37 °C incubator overnight, with agitation at 200 RPM. A 1 mL sample was then re-inoculated into 50 mL fresh LB and grown for approximately 2 hours to ensure cells were in exponential phase (final OD of 0.2 to 0.5) resulting in the starting population.

Devices were assembled for experiments by attaching outlet tubing to 60 mL syringes via needles, sealing channels with the agar sheet and placing the device in a protective casing. Growth medium was added to each reservoir. The starting population was placed onto the agar slab by streaking directly over the first channel using a sterile cotton swab. Finally, the device was sealed and incubated at 37 °C. To control humidity and avoid condensation at the lid, the lid of a commercially-available humidity chamber (ibidi, set to 90 % humidity) was placed over the device.

The outlet syringes for the channels were connected to a dual-syringe pump [32] set to apply suction at 0.75 mL/hr, with a 3 minute flush of 10 mL/hr every 3 hours to clear the flow of any air or solids. For antibiotic-free experiments, two channels were active at any one time: the seeded channel (C1) and adjacent channel (C2) were active at the experiment outset, and the next adjacent channel (C3) was activated and C1 was deactivated once the bacterial population approached C2. For antibiotic gradient experiments, four channels (C1, C2, C3, C4) were active through the duration of the experiment. The experiment was monitored until the bacterial population reached the third active channel, or a failure occurred. At this point the pump was stopped and the device disassembled and re-sterilised.

### Imaging and analysis

A Raspberry Pi Camera V2, held on a stand 30 cm above the device and controlled by a Raspberry Pi 4, was used to image the device from above at 30 minute intervals. A diffuse light source placed under the device provided backlight imaging. External light was blocked by an enclosure.

Image analysis to detect the bacterial front at each time point was achieved by applying background removal to images, followed by denoising and thresholding to separate opaque (microbial population blocking backlight illumination) from transparent areas (free agar). Background removal was performed by subtracting the first image from all subsequent images, thereby removing the channels and agar imperfections from the image. The de-noising process involved applying a Gaussian filter of 2 px standard deviation for each image of the stack as previously described [33]. To smooth variation in light availability between images, a Gaussian filter with standard deviation of 2 was performed between images, along the time axis.

To identify how far the population has grown, we used binary thresholding. To calculate the threshold value best fitting the data, we chose the Li algorithm as described in Ref. [34] as it best fitted manual thresholds chosen by eye. Areas identified by thresholding that were not contiguous to the core bacterial population were disregarded in the downstream analysis.

The bacterial front was quantified as the edge of the area identified as the bacterial population. With the image aligned parallel to the first channel, *p*_*i,t*_ denotes the boundary of the bacterial mass in either direction vertical to the channel, for each time *t* and position *i* along the streak. Using the length of a channel or diameter of the Petri dish as a guide allowed us to relate pixel to *µ*m on the device: approx. 730 *µ*m per pixel, apart from early control experiments at approx. 610 *µ*m per pixel.

We found heuristically that after 12 hours of incubation time the bacterial coverage was consistently dense enough to perform the image analysis outlined. The 12 hours time point therefore served as the initial time point for quantification of microbial growth, to better compare between experiments.

The average displacement, 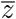, at time *t* was defined as the mean over the entire length of the streak, *L*, where the sum runs over individual pixels:

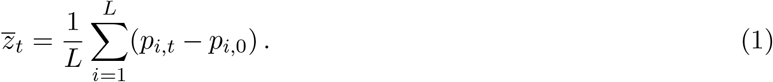

The mean speed of movement of the front, 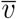, was calculated as the change in the mean displacement over time:

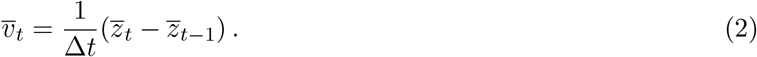

The roughness of the bacterial front, *r*, at each time point was calculated as the standard deviation of the front positions compared to positions in a smoothed contour of the front, *S*, using a Gaussian filter with a standard deviation of 7 mm:

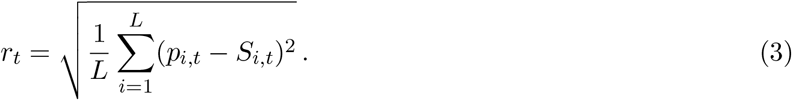

The inlet-outlet growth, Δ*v*, was calculated as the difference between the front speed at the first and last quarters of the population, corresponding to the inlet and outlet-side of the channel:

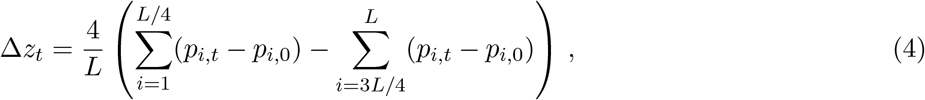

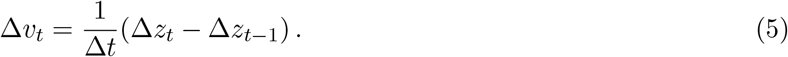

Front speed and inlet-outlet growth were plotted as a moving average (period of 6 frames) for clarity.This analysis was implemented in a Python script.

### Pressure limit analysis

Pressure limit testing was carried out experimentally concurrently with device development, by adjusting flow rate and device elevation. In addition, it is possible to model the pressure along the fluid flow as follows.

The device is composed of long, fixed-diameter tubes or channels connected together with varying elevations, so pressure was calculated following the integrated Poiseuille and Bernoulli equation as described in Synolakis & Badeer [35]:

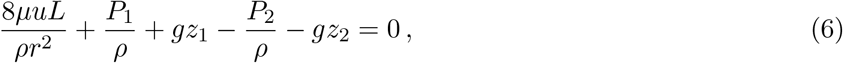

where indices 1 and 2 specify two points along a tube with radius *r*, separated by distance *L. P*_1_ and *P*_2_ are pressures at those points, *z*_1_ and *z*_2_ elevations at those points and *g* is acceleration due to gravity. Fluid viscosity (*µ*) and density (*ρ*) are assumed to be that of water at 37°C from literature [36].

A standard Bernoulli equation neglecting gravitational effects was used to describe pressure change at junctions, where there is significant sudden change in channel diameter:

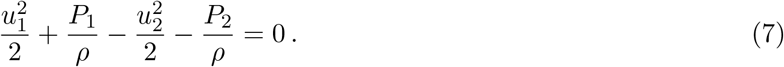

These equations were implemented in a Python script tracking pressure at key points within the device and periphery, starting from the pressure-equalised end of the set-up.

### Numerical simulation

Modelling of nutrient transport and mechanical properties of the device were carried out using finite element analysis (FEA) using COMSOL Multiphysics (COMSOL).

The nutrient transport model is a 3-dimensional model. It encompasses two channels with nutrient flow at a constant rate and with a steady-state laminar flow pattern from the inlet to the outlet. Above the channels, there is an agar sheet, modeled as a region of media with zero convective movement. Diffusion of nutrients is possible throughout the entire system, with an assumed diffusion coefficient of 6 *·* 10^−6^*cm*^2^*/s* in channels and in agar, taking glucose as an example for a molecule providing nutrients [5]. To one side of the channels, the agar extends to simulate a region without channels (channels without flow in experiments). To the other side, a reflecting boundary simulates a continuation of active channels. Diffusion through the device was modelled over 24 hours of simulated time.

The mechanical model describing deformation of the agar sheet due to negative pressure in the channel, was restricted to two dimensions since the device is invariant along the channel length. The model contains the polymer supporting structure, assumed not to exhibit deformation, and an agar sheet on top, overhanging the support. Symmetry constraints on either side simulate a continuation of channels. The Young modulus and Poisson ratio of 1.5% agar were assumed to be 52 kPa and 0.32 respectively [37]. Weight of the agar sheet was added to simulate gravitational effects, with density assumed to be that of water, and a pressure was applied to the overhanging agar to simulate the channel pressure (as calculated in section *Pressure limit analysis*). Deformation was calculated at steady state (Fig. S8).

Model accuracy was tested by decreasing mesh sizes until there were no visible differences in observables.

## Results

### Device design

We aimed to design a device that allows for continuous growth of microbes on an agar surface. To provide nutrients to replace those consumed by growing microbes and remove metabolic byproducts, the agar-based fluidic device needed a constant inflow and outflow of fresh and spent media, in analogy to microfluidic devices [11, 38] and chemostats [27] in micro-scale and planktonic culture methods, respectively. To maintain spatial structure on the agar surface, the flow of media was to occur in channels below the agar surface with exchange of media facilitated through a layer of agar via diffusion (Fig. 1A). To create a design which could be realised with resources accessible to microbiology laboratories, we opted for a core design on an agar slab on a polydimethylsiloxane (PDMS) base, shaped by 3D-printed template, connected to pumps and reservoirs using standard consumables (i.e. needles, syringes, tubing), see *Materials and Methods*.

Based on these design principles, we developed the device using a combination of experiments and modelling, in particular finite-element analysis (FEA) to interrogate nutrient transport and agar sheet stability. We based the overall device dimensions on a standard 96-well plate and chose an agar sheet height of 1.7 mm that allowed to achieve a nutrient concentration at the agar surface sufficient to sustain microbial growth (Fig. S1A) while ensuring mechanical stability (Fig. S8). As mentioned, the supporting structure allowing media flow was made of PDMS, a transparent and biocompatible material [39] that allows recessed parallel channels for media flow to be patterned via soft lithography (Fig. 1). We found that a channel width and height of 1 mm and a separation size between channels of 5 mm provided a uniform nutrient concentration on the agar surface (Fig. S1) and good agar adhesion (Fig. S8).

To ensure stability of the device, we aimed for negative pressure within the device, pulling the agar sheet towards the base, achieved with syringe pumps connected to each channel outlet and media pulled through each channel independently (Fig. 1A). Modelling showed the pressure within channels relative to ambient to depend linearly on the medium flow rate (Fig. S7) and with the agar sheet becoming deformed when channel pressure was more than 1 kPa. Experimentally, we found that a flow rate of 0.75 mL/hr with a burst of 10 mL/hr flow for 3 minutes every 2 hours successfully kept the channels clear of air and sustained long-term and homogeneous growth. Note that for these parameters, flow of growth medium through the device is laminar (i.e., with a maximum Reynold’s number of 3 at the maximum flow rate of 10 mL/hr).

Modelling showed that for these optimised parameters, the nutrient profile at the agar surface quickly approaches stability. The nutrient concentration close to channels reaches 80% of that in media within 12 hours, and the difference in nutrient concentration between the channel inlets and outlets sides of the device is negligible by 24 hours (Fig. 2).

**Figure 2:**
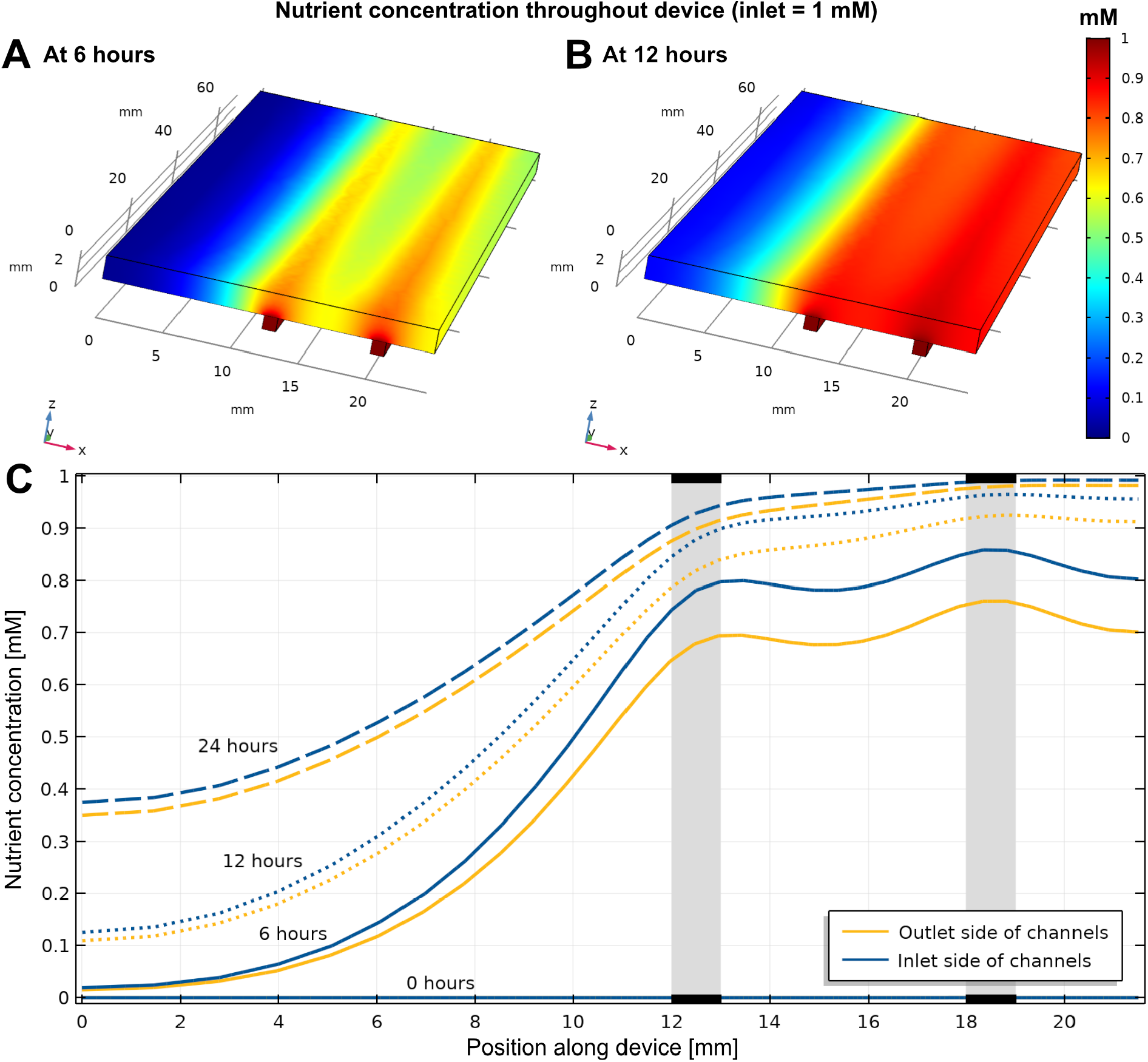
Simulated nutrient concentration profile in the device. (A-B) Three-dimensional representation of the agar sheet with two channels underneath supplying nutrients at 6 and 12 hours after the start of supplying nutrients through the channels with no nutrients present in the agar sheet initially. Simulation has been performed using COMSOL using device parameters described in Table S1. (C) Surface concentration profile of nutrients along the device, showing the profile at the boundary associated with inlets (blue) and outlets (yellow) at 0, 6, 12 and 24 hours. Locations of active channels are indicated in grey.

To allow for quantitative characterization of microbial growth and expansion in the device, we illuminated the device from below (back-light illumination), recorded images using a low-cost camera and single-board computer, and performed quantitative image anaysis, see *Materials and Methods*.

### Sustained growth and expansion of *E. coli* populations in the agar-based fluidic device

To demonstrate the ability of the device to continuously supply nutrients to a growing microbial population for an extended period of time, we investigated the sustained growth of a population of the bacterium *Escherichia coli*, more specifically the non-motile (on hard agar) strain DH5*α*. We seeded *E. coli* populations as streaks on the agar surface of the device, of approximately 1.5 mm width and 70 mm length (Fig. 3A) with the agar sheet initially devoid of nutrients. Bacteria were inoculated above a channel with continuous nutrient flow (C1), dubbed ‘active’ in the following; one of the neighbouring channels (C2) was also active, while the channel on the other side (C0) remained empty (top and bottom channels in Fig. 3A, respectively). Once the bacterial front approached C2, the adjacent channel (C3, not shown) was activated and C1 was deactivated.

**Figure 3:**
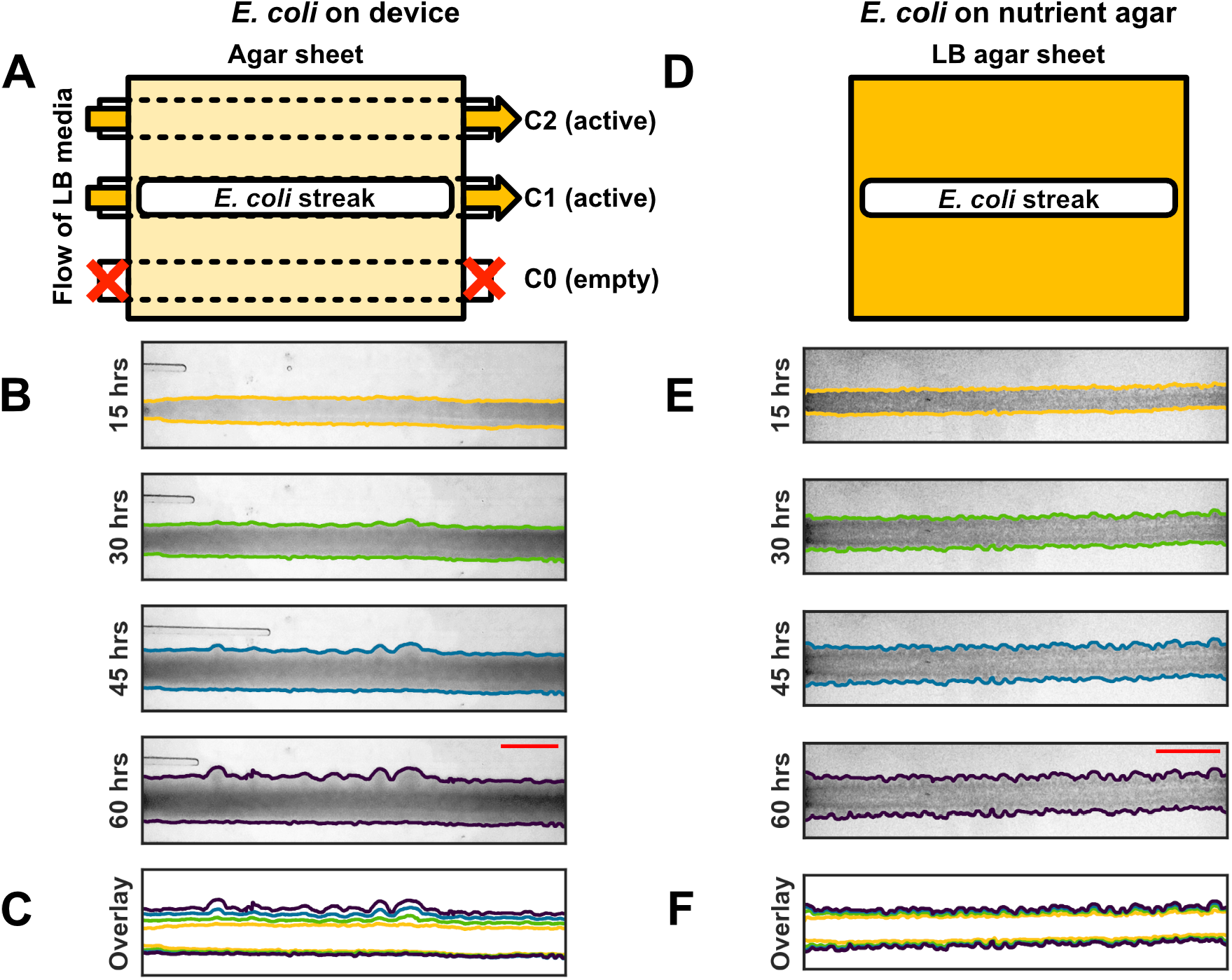
Experimental design and spatial expansion of *E. coli* population. (A)Diagram of the device from above with two adjacent active channels and one empty channel as well as a streak of *E. coli* over the middle channel. (B) Images of the population (dark region) taken from above the device with illumination from the bottom, in the same orientation as panel (A) with fronts identified by image analysis overlayed as coloured lines. (C) Merged overlay of tracked population fronts to visualise propagation of the front over time, colours correspond to the colours and thus time points in panel B. (D-F) Corresponding population expansion on a an agar sheet with nutrients but without flow below. Scale bars represent 1 cm. For complete time lapse information see Videos S1,S2.

Populations were not identifiable immediately after inoculation due to the initially low bacterial density, similar to what observed on standard agar plates. From 12 hours after seeding, bacterial density was sufficient to allow for quantitative image analysis (*Materials and Methods*) to identify the spatial boundaries of the population as overlaid onto recorded images in Fig. 3B.

Fronts at different times are displayed together in Fig. 3C. While boundaries are clearly separated on the side of the active channel, they are much closer to each other on the side of the empty channel, illustrating that as time passed the bacterial population expanded mainly towards the active channel. In contrast, the same strain of *E. coli* ceased to grow across the standard agar plate after 45 hours (Fig. 3D-F). In both setups, the front is undulating as viewed from left to right; corresponding to tiny pockets of bacteria on the nutrient agar plate, and as larger and more sporadic pockets for the population on our new device.

Following identification of the fronts, we extracted quantitative measurements from the images above by assigning each bacterial front a mean displacement 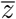 (Fig. 4A), mean speed 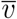 (Fig. 4B), and roughness *ρ* (Fig. 4C), see *Materials and methods* for details. Note that roughness indicates how undulating a front is compared to its smoothed contour, but does not contain information on spatial correlation, i.e., whether the pockets of bacteria are large and sparse or dense and small (as seen in Fig. 3F and C, respectively).

**Figure 4:**
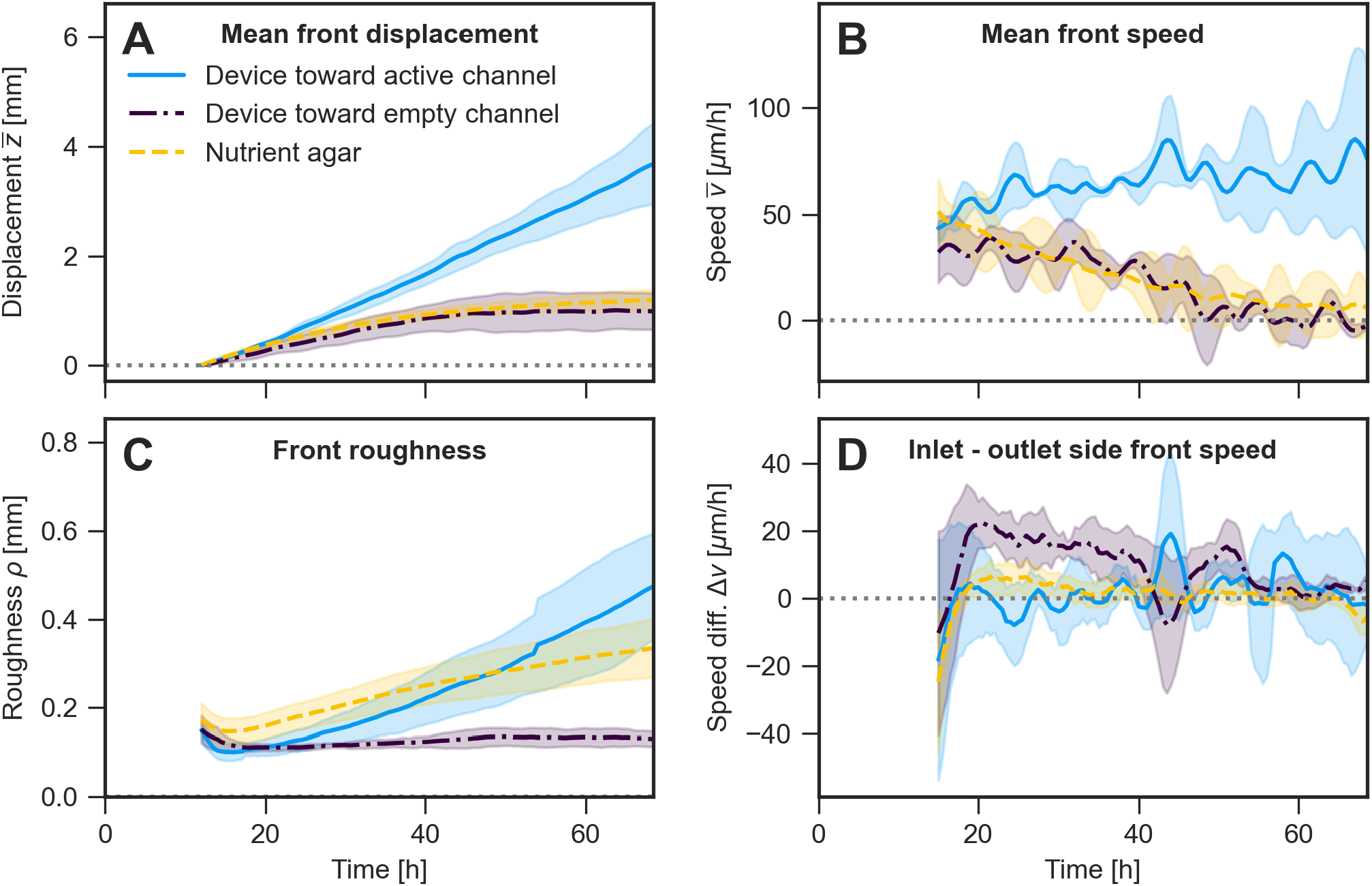
Characterisation of movement of *E. coli* population front. (A)Mean displacement of the bacterial front from the initial position over time on the agar-based fluidic device, towards the active channel (blue solid line) and towards the empty channel (purple dash-dotted line), as well as in both directions on a standard agar plate (yellow dashed line). The mean and standard deviation of three independent experiments are displayed as lines and shaded areas, respectively. (B) Mean speed of the bacterial front over time, calculated as a moving average over 6 time points. (C) Roughness of the front over time, calculated using the standard deviation of the front from its smoothed contour. (D) Difference in front speed over time at the device boundaries associated with inlets and outlets, respectively. The mean and standard deviation error of three independent experiments are displayed as lines and shaded areas, respectively.

We found that the lateral growth of *E. coli* towards an active channel was sustained for the 70 hour duration of the experiment, with the average population front advancing about three times further towards the active channel than either towards the empty channel or on the control plate (Fig. 4A). Front propagation on a standard agar plate began with the same speed as on the device, about 50 µm/h, but decelerated steadily, approaching zero towards the end of the experiment’s duration. This is also the case for the front growing towards the empty channel in our device, indicating a gradual shift in both cases towards halted growth of bacteria at the front (Fig. 4B).

Roughness of the front growing towards the active channel (Fig. 4C) is comparable to that on standard agar plates. The mean active channel front roughness is initially somewhat smaller than on the control nutrient agar plate, but increases over the course of the experiment, passing the control plate roughness by about 50 hours and reaching about 0.5 mm by the end of the experiment. Front roughness on the control plate also increases over time, but with only a two-fold increase from below 0.2 mm to about 0.3 mm over the course of the experiment. Population fronts moving towards an empty channel exhibit sustained low roughness throughout the experiment, at the same level as the active channel front at the start of the experiment.

We expected the medium composition to vary with the location along the channel due to nutrient exchange with the agar slab. These differences could cause variation in growth along the front. To investigate such an effect, we computed the difference Δ*v* between front speed at the inlet and outlet side of the channel (Fig. 4D). This speed difference is compatible with zero for the population moving towards an active channel; the control plate without channels shows a similar variation. We observe a difference for the population front moving towards the empty channel, indicating that the environmental heterogeneity across the length of the channel has an effect at the lower nutrient concentrations bacteria are subjected to.

### Sustained growth and expansion of motile bacteria in the agar-based fluidic device

To test our agar-based fluidic device further, we investigated growth of *Pseudomonas aeruginosa*, which is a motile and chemotactic bacterium as well as a model organism for studies of biofilm formation. We characterised growth of strain PA14 (parental strain, *PS*, in the following) and compared its growth to the growth of a flagella-knockout with reduced motility (PA14 *flgK::Tn5B30(Tc*^*R*^), Δ*flgK* in the following).

The population front of the parental strain *P. aeruginosa PS* propagates fast across the surface (Fig. 5A,B) with a significant slowdown within the first 30 hours (Fig. 5F,G). The Δ*flgK* population displays linear expansion towards the active channel (Fig. 5C,D) with pattern and speed similar to what we observed earlier for *E. coli* (Fig. 5F). The heterogeneity in population growth in most experiments carried out with strain *PS* (Fig. 5B) resulted in a smooth population front with more extensive growth at the channel inlets than the channel outlets, a trend visible to a much lesser extent for the Δ*flgK* populations (Figs. 5D and I). Front roughness is similar across all populations (Fig. 5H), with the exception of one PS experiment that had a higher roughness (individual experiments shown in Fig. S3) affecting average roughness.

**Figure 5:**
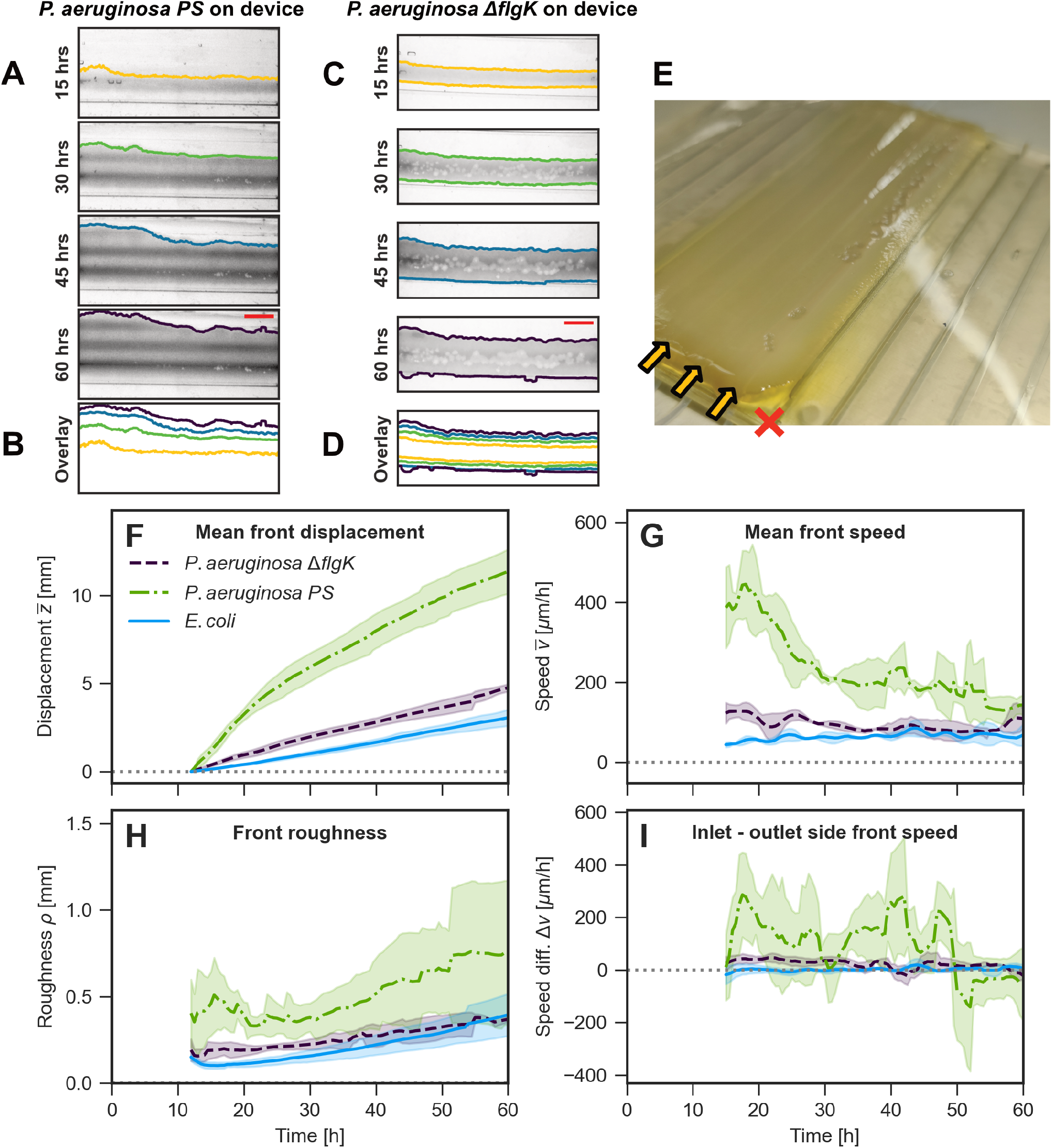
Characterisation of movement of *P. aeruginosa* population fronts and comparison to that of *E. coli*. (A) Images of a population of *P. aeruginosa PS* growing in the device, analogous to Fig. 3B. Scale bars represent 1 cm. (B) Population fronts moving towards active channel merged as in Fig. 3C. (C-D) Like (A-B) for a population of *P. aeruginosa* Δ*flgK*. (E) Photograph of *P. aeruginosa PS* population on agar sheet around the channel that remains empty (marked by red cross, opposite to the front growing towards the active channel). Note the striated patterning of the front as well as clearance plaques. (F-I) Mean front displacement, mean front speed, front roughness, and difference in speed between the inlet and outlet sides, as in Fig. 4, for both strains of *P. aeruginosa* and compared to *E. coli*. For complete time lapse information see Videos S3,S4. The mean and standard deviation error of three independent experiments are displayed as lines and shaded areas, respectively.

We observed significant difference in expansion towards the empty channel when comparing *PS* and Δ*flgK* strains. The *PS* populations spread outwards, quickly propagating towards the empty channel, but with very low population density. This density decreases the further cells are from the nutrient source, forming a gradient stretching away from the initial population (Fig. 5E). This observation was consistent throughout experiments, but could not be quantified with the image analysis used. The Δ*flgK* strain, in contrast, consistently formed a visible and demarcated population front, similar to *E. coli*. Unlike *E. coli*, however, the populations visibly grew towards the empty channel (Fig. 5D) for the first 45 hours, seemingly less affected by the low nutrient availability.

While these experiments were designed as proof of principle for bacterial growth well beyond the duration of growth on agar slabs in Petri dishes, comparing growth on agar vs our new device revealed differences emphasising the role even subtle environmental conditions play in microbial colony growth. First, while the faster speed of the parental strain compared to the flagella knockout Δ*flgK* on the device is at first not surprising, both strains showed similar growth patterns on standard agar plates (Fig. S4). Second, all experiments performed with either strain of *P. aeruginosa* in the device resulted in plaque-like clearings in the initial seeded area (Figs. 5C,E).

### Growth and expansion of *P. aeruginosa* in a spatial antibiotic gradient

The device does not only offer a continuous supply of nutrients and removal of metabolic byproducts. By providing locations (areas around channels) with fixed media composition, it also allows one to create temporarily stable gradients. To demonstrate this capability, we simulated antibiotic gradients originating solely from fixed concentrations at individual channels and subsequently investigated *P. aeruginosa*’s growth and expansion in such gradients.

Solving the diffusion equation predicts a stable gradient to emerge between two regions at fixed antibiotic concentration. To confirm this prediction for channels with an agar sheet on top, we again modelled transport and diffusion (in the absence of bacteria) using finite element analysis with media flowing through channels, now with different concentrations of an antibiotic (Fig. 6). Starting with an agar sheet without antibiotic, modelling predicts the gradient to emerge rapidly and to be stable by 12 hours, the point when bacterial populations became visible in our experiments. Due to finite thickness of the agar sheet, the antibiotic concentration at the agar surface is not equal to the concentration in the channel; in particular we predict a small concentration of antibiotic even above the channel when the antibiotic is not pulled through the device (leftmost channel in Fig. 6). We realised the scenarios simulated experimentally, by pulling medium through channels with specified concentrations of the cephalosporin antibiotic ceftazidime [40], known to act against *P. aeruginosa*, and asked how bacterial populations grow and expand in the presence of these gradients (Fig. 7).

**Figure 6:**
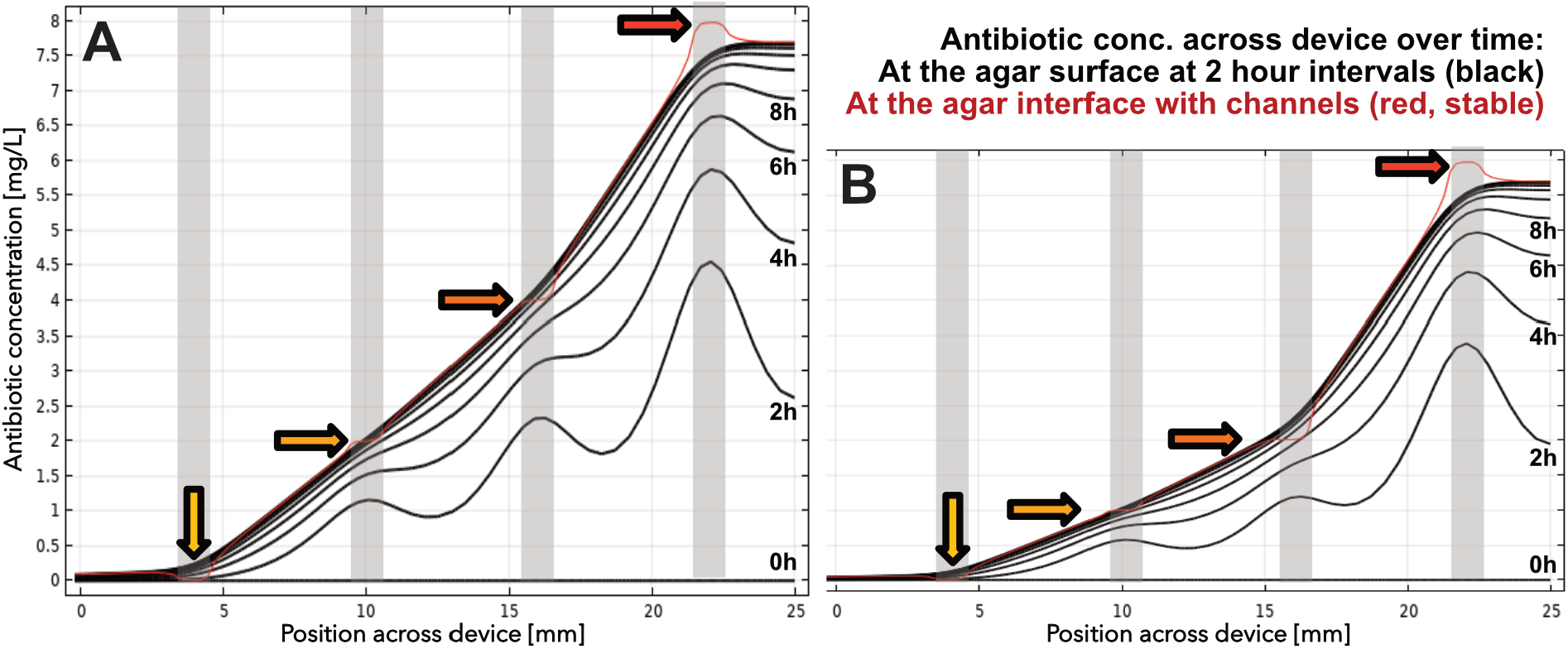
Simulated emergence of an antibiotic gradient in the device. Development of a (A) steep and (B) shallow gradient across the agar surface at two hour intervals in black, with the corresponding concentration at the base of the agar sheet adjacent to channels marked in red. Empty channel to the left of the antibiotic-free channel is not shown. Antibiotic concentration at each media channel (location marked in grey) is shown with an arrow.

**Figure 7:**
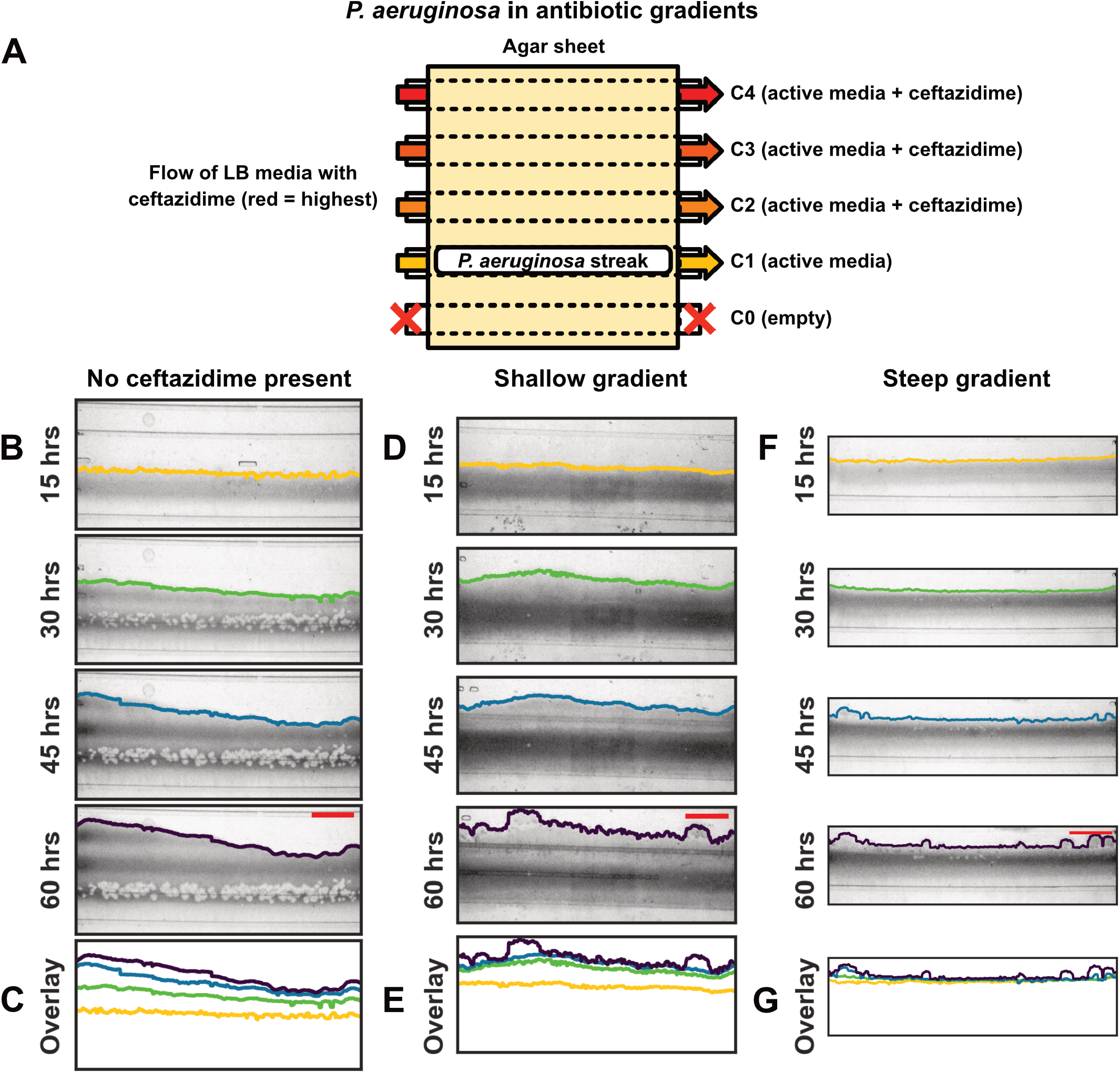
Front development of *P. aeruginosa* populations by antibiotic gradient. (A) Diagram of the device from above with four adjacent active channels (with increasing concentration of ceftazidime, indicated by colour) and one empty channel as well as a streak of *P. aeruginosa* over the active channel without antibiotics. (B,D,F) Images of a population (dark region) on the device in the same orientation as in panel (A) for the control (no gradient), a shallow (to 3/4× MIC) and a steep (to 1× MIC) gradient with fronts identified as coloured lines. Scale bars represent 1 cm. (C,E,G) Merged overlay of tracked population fronts.

*P. aeruginosa* populations were deposited as streaks on the agar surface above a channel with constant nutrient flow and no antibiotics. On one side, three neighbouring media channels were permeated with either 1, 2 and 6 mg/L of ceftazidime (1/8×, 1/4× and 3/4× the minimum inhibitory concentration (MIC) value measured against *P. aeruginosa*), forming a shallow gradient, or 2, 4 and 8 mg/L ceftazidime (1/4×, 1/2× and 1× MIC), forming a steep gradient (Fig. 7A). On the other side, a channel was kept empty, towards which we expected no or minimal growth based on the experiments without antibiotics (Fig. 7A). Growth of bacterial populations was followed and analysed as described above (Fig. 7B-G) together with experiments in the absence of antibiotic gradients.

We found that the front propagation along the gradient anticorrelated with the steepness of the gradient (7B-G). This anticorrelation was expected, given that the steeper gradient could (i) lead to cells dividing more slowly, (ii) lead to the evolution of new phenotypes with a trade-off in growth rate and antibiotic tolerance, (iii) increase the time taken for cells to adapt successfully to the antibiotic concentration or (iv) provoke a negative response in the chemotactic population, leading to movement away from the gradient. Quantitative analysis revealed that growth in the presence of a shallow gradient, while slower, is similar to the ‘no gradient’ scenario, as the population initially expands quickly before slowing down. The steep gradient, however, suppressed population growth until about 30 hours after which the front propagated (Fig. 8A-B).

**Figure 8:**
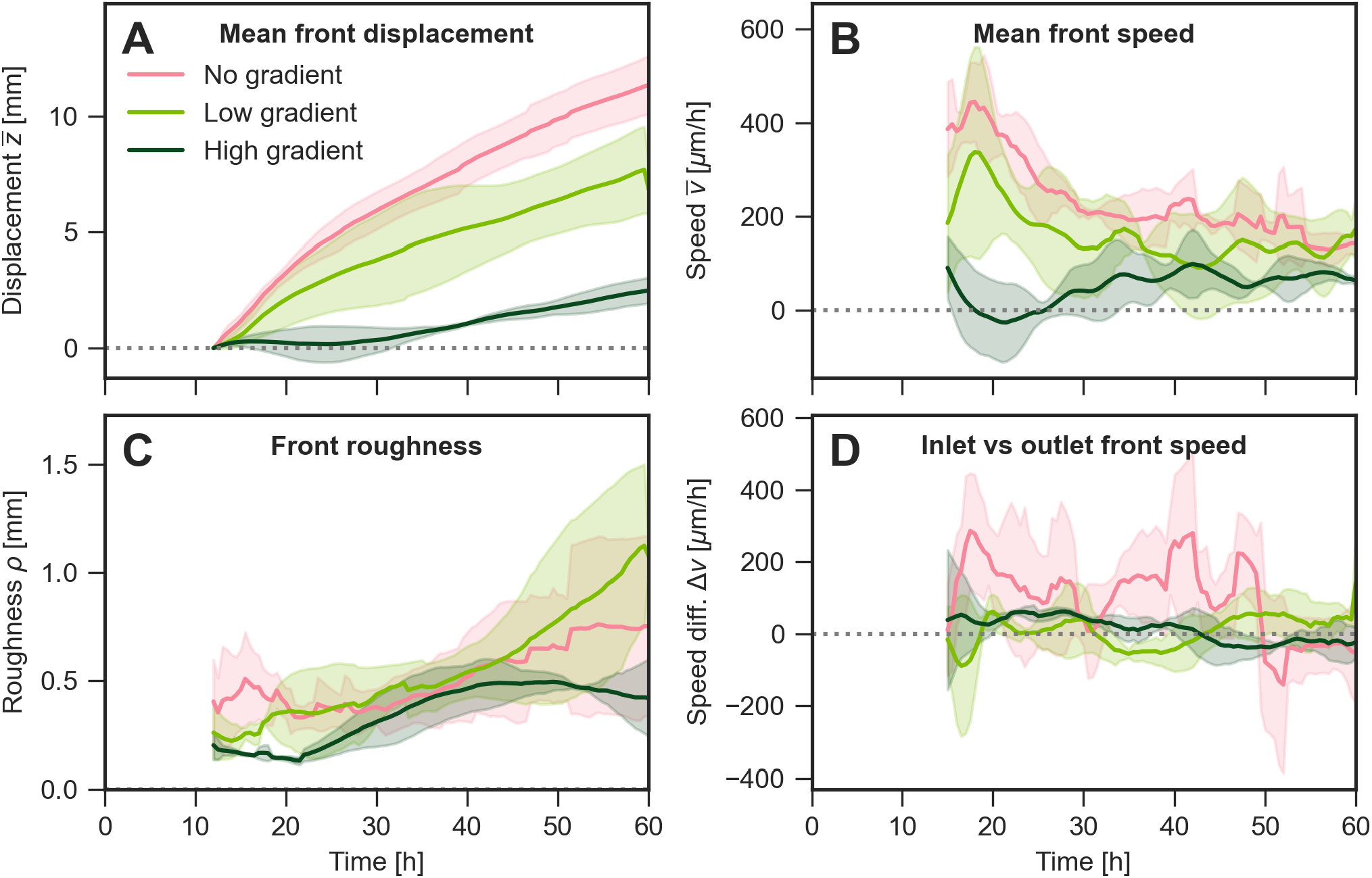
Analysis of bacterial front movement by gradient type for *P. aeruginosa PS* populations. (A)Mean front displacement, (B) mean front speed, (C) front roughness, and (D) difference in speed between the inlet and outlet sides, as in Fig. 4, with movement towards an antibiotic-free active channel (pink line), a shallow gradient of 0 to 1/4x MIC over three channels (light green line), and a steep gradient of 0 to 1/2x MIC over three channels (dark green line) and an empty channel behind the population. The mean and standard deviation error of three independent experiments are displayed as lines and shaded areas, respectively.

In the presence of a gradient, protrusions in the front emerged in the second half of the experiment (Figs. 7D-G). They appear qualitatively different to the broader variation observed in the absence of a gradient (Figs. 7B,C).

## Discussion

Culturing microbes on agar is an integral part of microbiological history and practice, but the local environment cannot be quantitatively controlled, neither temporally nor spatially, due to non-directional molecular diffusion within the agar. Tight temporal and spatial environmental control can be achieved in microfluidic devices, for example by growing cells in a gel or between agar and a harder material with media exchange provided in close proximity [41–45]. However, these devices are limited to the study of only a small number of cells in great detail instead of microbial populations. Furthermore, cells are highly constrained and sealed off from the gaseous environment, preventing the growth of three-dimensional structures and access to the gaseous environment, which can play an important role in the development of microbial communities [46].

To overcome these limitations while maintaining most of the benefits of agar cultures, we developed a millifluidic device that allows users to tightly control the liquid environment under the agar sheet. The device allowed for sustained growth of *E. coli* and *P. aeruginosa* populations at the agar-air interface for at least 60 hours. Surprisingly, we observed qualitative differences in *P. aeruginosa* colony growth between our new agar-based milli-fluidic device and conventional agar plates. First, we noticed a difference in colony growth between *P. aeruginosa* Δ*flgK* and its parental strain that is absent on control plates (Figs. S4 and S5). Both strains of *P. aeruginosa* used are capable of twitching motility via pili on a hard surface [29, 47], whereas only the parental strain is capable of swarming motility due to the presence of flagella. On our milli-fluidic device, the parental strain propagated much faster than the flagella-knockout (Fig. S4B), hinting to a role of flagella-mediated motility in the population expansion that is absent on passive agar plates. Further research is necessary to elucidate whether this is a consequence of differences in cell states between the device and control plate or a result of potential difference in the physico-chemical properties of the agar surface such as wetness of the surface. In either case, this observation highlights the importance of the environment on microbial growth and calls for both further quantitative studies and serves as a caution against generalising findings from growth on traditional agar plates.

In addition, we consistently observed clearance plaques on the device within the area that *P. aeruginosa* populations were seeded in. These were absent on control plates. This may be attributable to over-expression of the *Pseudomonas* quinolone signal (PQS), which is rarely visible in populations grown on nutrient agar but is thought to be displayed within highly dense populations [48]. The constant provision of nutrients in the device should result in higher density growth than on nutrient agar, making over-expression of PQS a viable mechanism to describe the appearance of the clearance plaques. Future research could utilize the temporal control the device provides to further investigate this phenomenon.

Our new agar-based millifluidic device is significantly different from the metre-scale MEGAplate where chemotaxing bacteria expanded against a step-wise antibiotic gradient [8]. To achieve a stable gradient at the time scale of the experiment, the authors used a very large plate, which in turn requires active motility for the population to expand over those distances. The device presented here, however, can provide a stable gradient on a much smaller scale and thus allow to study bacterial expansion without active dispersal.

Moreover, the MEGAplate [8] used antibiotic steps rather than a linear gradient employed here and focused on concentrations orders of higher than the MIC while we focused on sub-MIC concentrations. An intriguing observation obtained via the MEGAplate was the growth patterns forming when reaching the next higher antibiotic concentration, with population growth en-masse halting at the boundary and then continuing from adapted individual bacteria at the population front, spreading out into the available space [8]. Despite the continuous gradient in our device, there was similarly significant bulging of the population from individual locations across the initial seeded population (Fig. 7C, E and G). This heterogeneous front then remained for the remainder of the experiment in the milli-fluidic device. The bacteria were therefore either not able to adapt continuously to the linear gradient, or gained a higher level of resistance than required when an adaptation occurred.

The overall front propagation towards the gradient was slower when the gradient was steeper both in our device and in Baym *et al*. [8]. This is in contrast with Lapińska *et al*. [14], where fast growing phenotypic variants in single-cell environments were shown to avoid accumulation of a different antibiotic, the macrolide roxythromycin, suggesting that the relationship between bacterial growth and antibiotic efficacy varies with the antibiotic in use.

In our device, there is significant variation in displacement and front speed difference between channel inlets and outlets as well as between replicates of no-gradient and shallow gradient experiments. However, there is very little variation at a steep gradient (Figs. 8A/D and S6A/D). Baym *et al*. [8] found a wider range of adaptations for bacteria encountering a smaller antibiotic step than a large one. This means the subpopulations that arise in these experiments are likely more heterogeneous in growth. We expect such heterogeneity to be reflected in heterogeneity of population expansion; a steep gradient leading to a more homogeneous population is therefore consistent with our results.

It is also possible that a gradient results in a negative response in a chemotaxing population, leading to movement away from the gradient and slowing propagation. Findings by Oliveira *et al*. [23] conclude that single cells of *P. aeruginosa* chemotax towards antibiotics and that no chemotaxis is visible at all at the population level. In our experiments, however, chemotaxis is seemingly at play in behaviour of the front moving towards empty channels in our device. Growth in this direction is not analysable quantitatively due to the population density gradient that forms for *P. aeruginosa* (Fig. 7); however, we observe qualitatively that there is stronger growth towards empty channels for shallow and steep gradients than in the absence of a gradient.

Moreover, while in Oliveira *et al*. [23], single cells of *P. aeruginosa* were shown to chemotax towards antibiotic gradients in an attempt to eradicate a threat, the study also found that these cells moving towards antibiotics are then unable to grow. Any growth or movement of single cells towards antibiotics in our experiments, where cells then die or remain inhibited, would not be visible using our analysis methods as we cannot pick up low densities of cells. This limitation highlights the importance of both single-cell and population level research into antibiotics and environmental variation; the addition of single-cell microscopy, or use of our device in conjunction with smaller microfluidic devices, would be necessary to understand the interplay between these two scales and will allow to achieve an more holistic understanding of bacterial behaviour.

We here focused on expansion of bacterial colonies for days and growth within antibiotic gradients. Changing the layout of the channels within the PDMS support structure could provide a myriad of different environments; changing the flow over time allows design of environments that are not only varying spatially but also temporally. As the core of the device can be manufactured with access to a 3D printer and low-cost materials, we hope that our device could be a starting point to explore bacterial growth and dispersal on complex, yet tightly controlled, surfaces.

## Acknowledgements

Brandon Tuck was supported by an EPSRC DTP PhD studentship EP/R513210/1. Stefano Pagliara was supported by a BBSRC research grant BB/V008021/1, and Wolfram Möbius acknowledges support by BBSRC via BBSRC-NSF/BIO grant BB/V011464/1. We are also grateful for input from Sean Booth on *P*.*aeruginosa* plaque formation.

## Supplementary Material

**Table S1:** Parameters used in device and COMSOL simulations.

|  |  |
| --- | --- |
| Channel height | 1 mm |
| Channel width | 1 mm |
| Channel length | 70 mm |
| Agar height | 1.7 mm |
| Channel separation | 5 mm |
| Diffusion coefficient | $6\text{E-}6 \text{ cm}^2/\text{s}$ |
| Simulated flow rate | 1 mL/hr |
| Flow rate | 0.75 mL/hr for 2 hrs / 10 mL/hr for 2 mins |
| Tube inner/outer diameter | 0.5 mm/1 mm |
| Inlet tube length | 45 cm |
| Outlet tube length | 30 cm |
| Tail length | 12 mm |
| Young modulus | 52 kPa |
| Poisson ratio | 0.32 |

**Figure S1:**
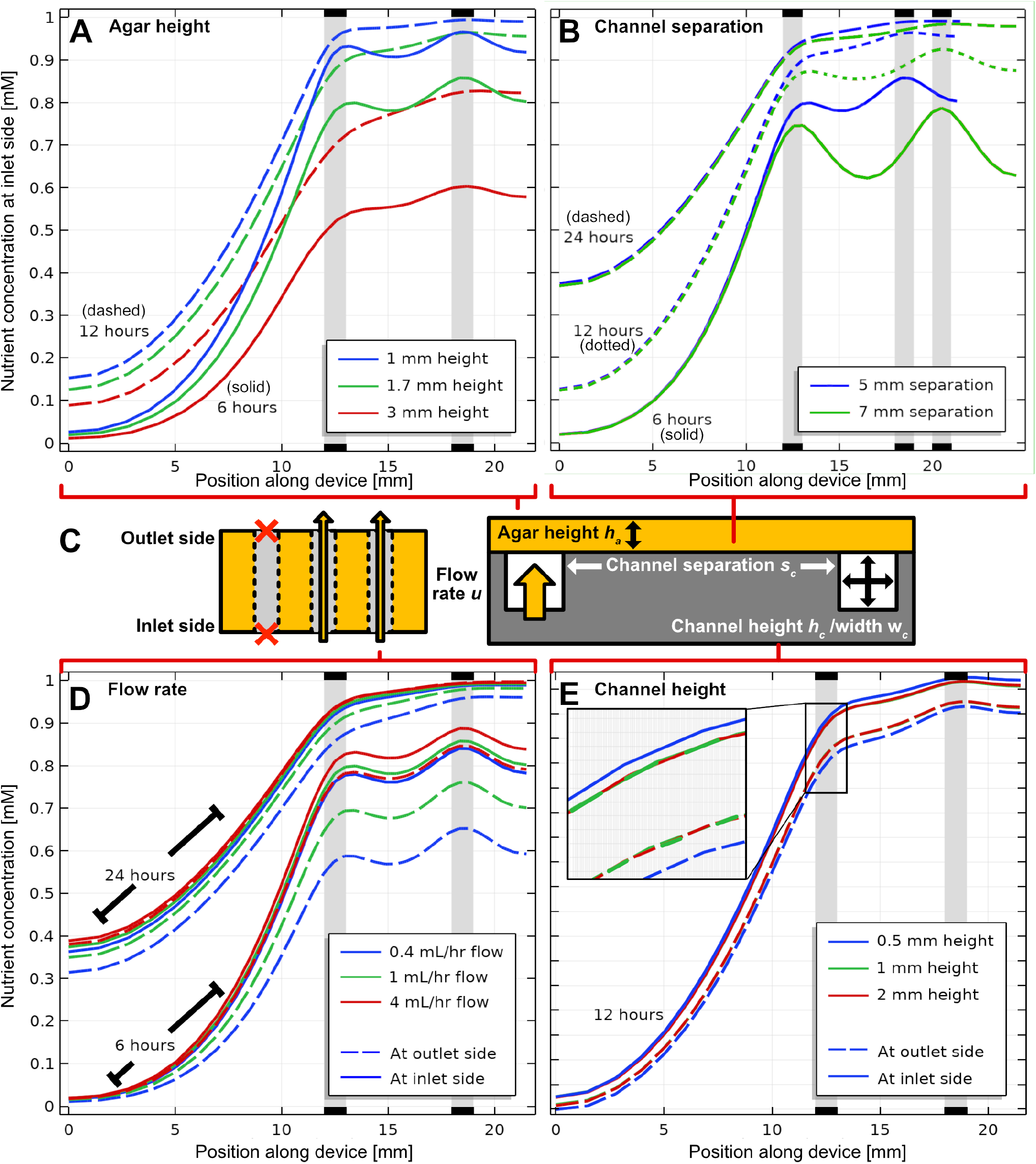
Simulated effect of varying device parameters on nutrient concentration on agar surface in absence of bacterial growth, by time since the start of the experiment. A) Agar height at 6 (solid) and 12 (dashed) hours. B) Channel separation at 6 (solid), 12 (dotted) and 24 (dashed) hours. C) Diagram of device from overhead (left) and side (right) showing parameters and location of inlet and outlets. D) Flow rate in channels at 6 (bottom) and 24 (top) hours, nutrient concentration at inlet and outlet side. E) Channel height at 12 hours, nutrient concentration at inlet and outlet side. Inset is area over the seeded channel showing location of 1 mm height which is not visible in the main figure. All non-specified parameters are listed in Table S1.

**Figure S2:**
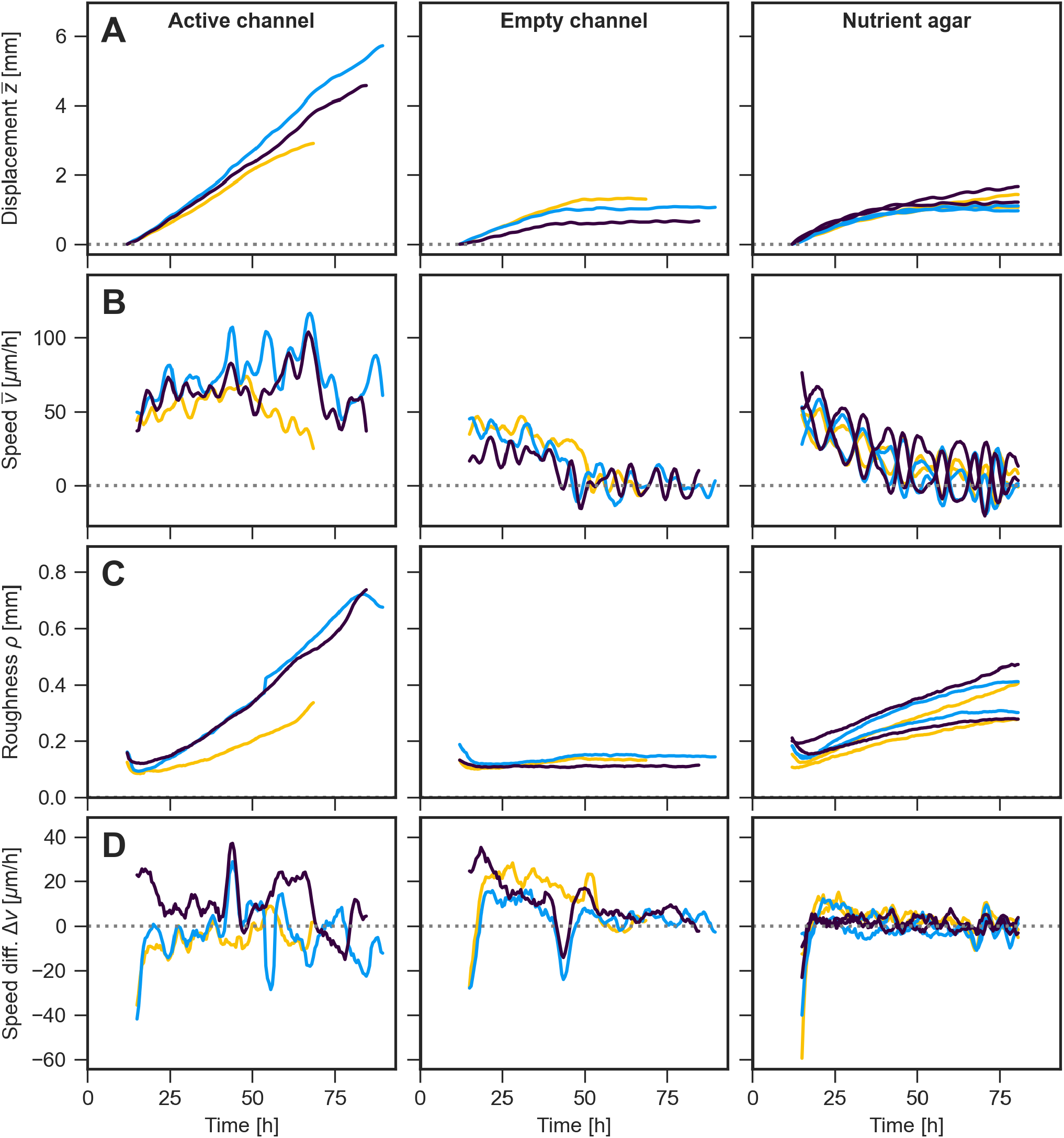
Data for individual replicates averaged in Fig. 4 on characterisation of movement of *E. coli* population fronts. A)Distance of the bacterial front from the initial position as averaged across the front over time on the agar-based fluidic device, towards the active channel (left) and towards the empty channel (middle), as well as on a standard agar plate (right). Individual replicate experiments are indicated in blue, purple, and yellow respectively, with two fronts for each experiment on a standard agar plate. B) Like panel A, but for mean speed of the bacterial front over time, calculated as a moving average of 6 time points over time. C) Like panel A, but for roughness of the front over time, calculated using the standard deviation of the front from its smooth contour. D) Like panel A, but for difference in front speed at the device boundaries associated with inlets and outlets, respectively.

**Figure S3:**
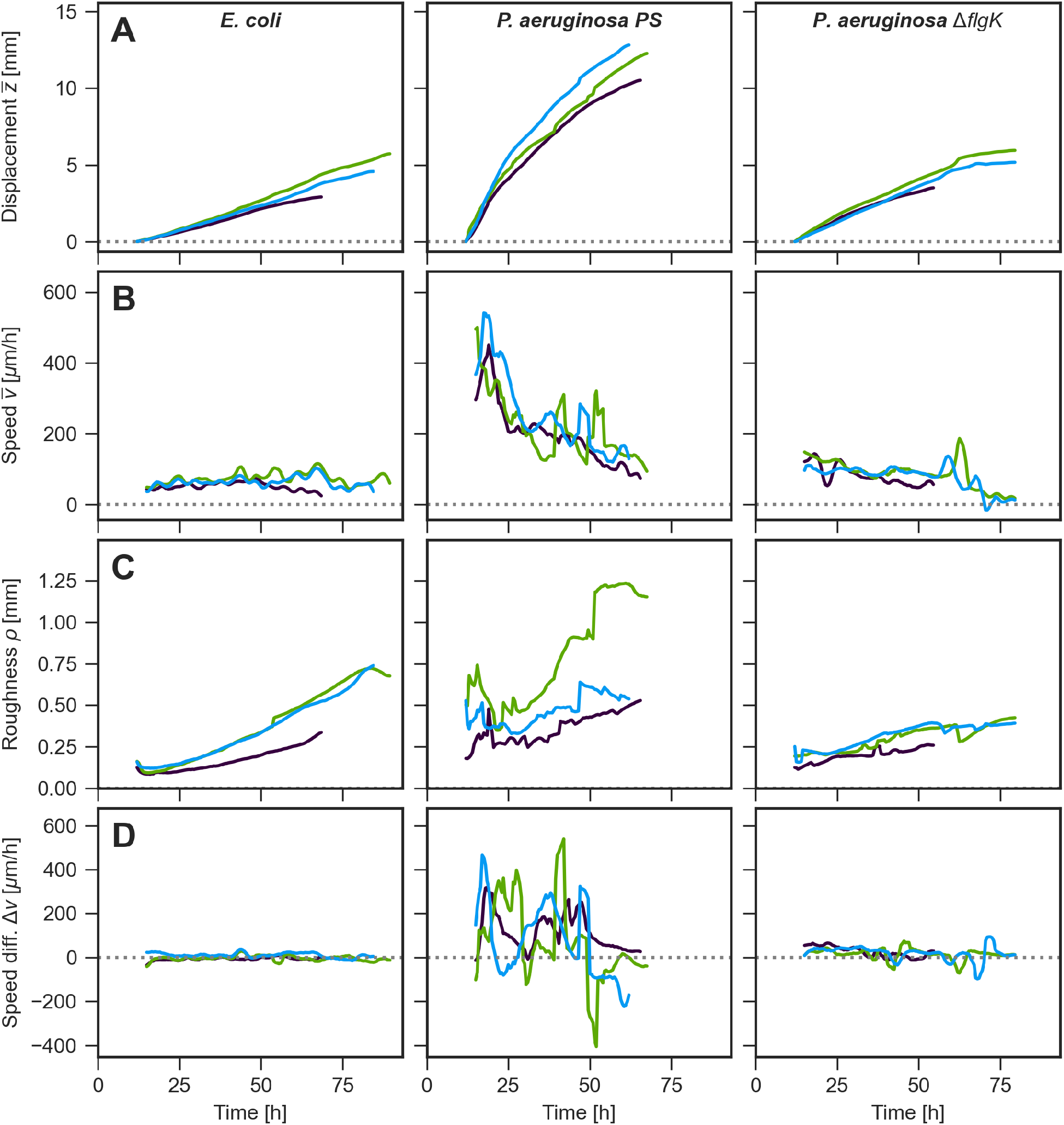
Data for individual replicates averaged in Fig. 5 on characterisation of movement of population fronts on the device toward an active channel. A) Distance of the bacterial front from the initial position as averaged across the front over time on the agar-based fluidic device, for *E. coli* (left) and *P. aeruginosa PS* (middle), as well as the Δ*flgK* variant of the latter (right). Individual replicate experiments are indicated in purple, green and blue. B) Like panel A, but for mean speed of the bacterial front over time, calculated as a moving average of 6 time points over time. C) Like panel A, but for roughness of the front over time, calculated using the standard deviation of the front from its smooth contour. D) Like panel A, but for difference in front speed at the device boundaries associated with inlets and outlets, respectively.

**Figure S4:**
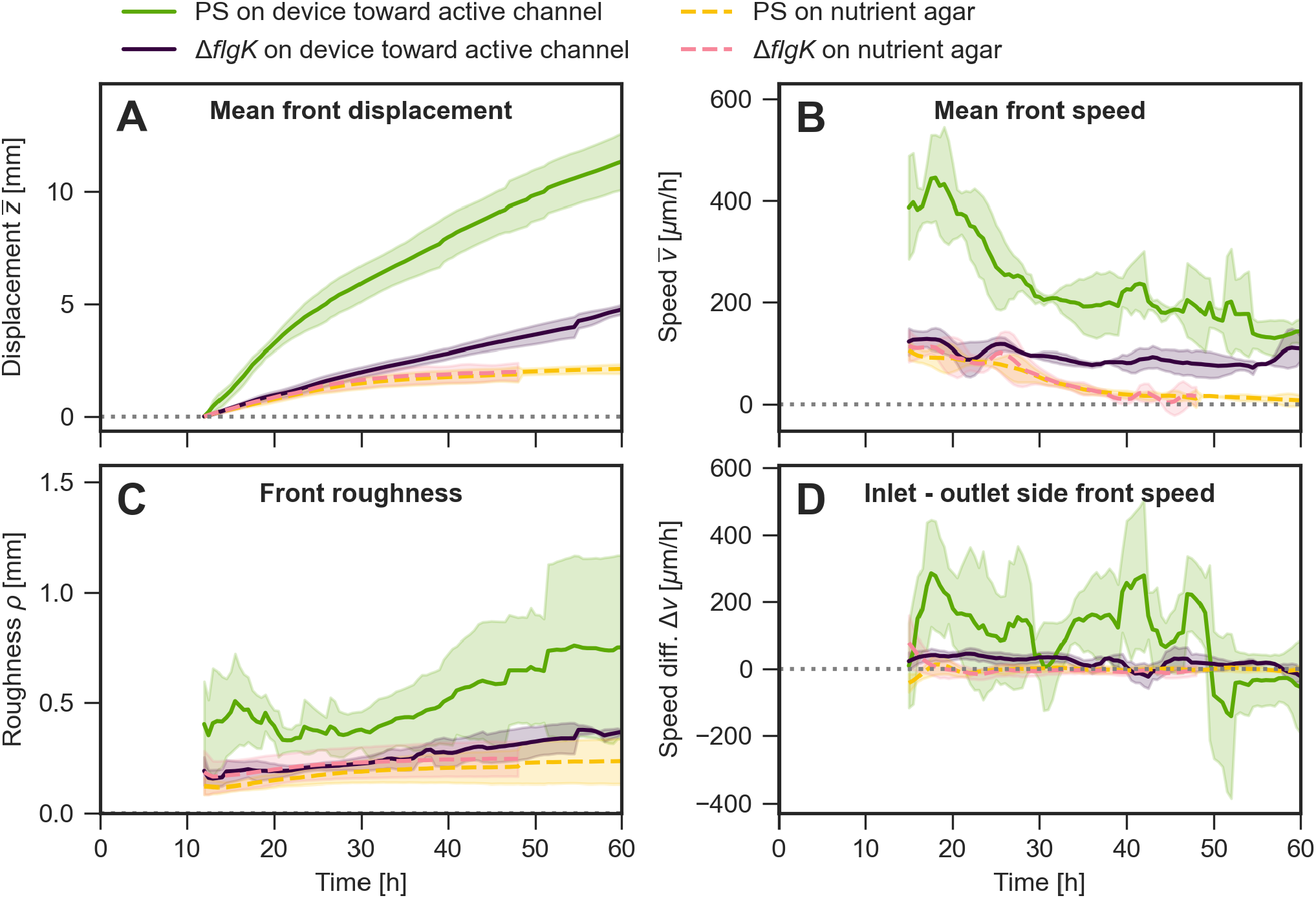
Characterisation of movement of *P. aeruginosa* population front. A-D) Displacement, speed, roughness and difference in speed between the inlet and outlet side, as in Fig. 4, for both strains of *P. aeruginosa*, on the device towards the active channel and also on standard nutrient agar.

**Figure S5:**
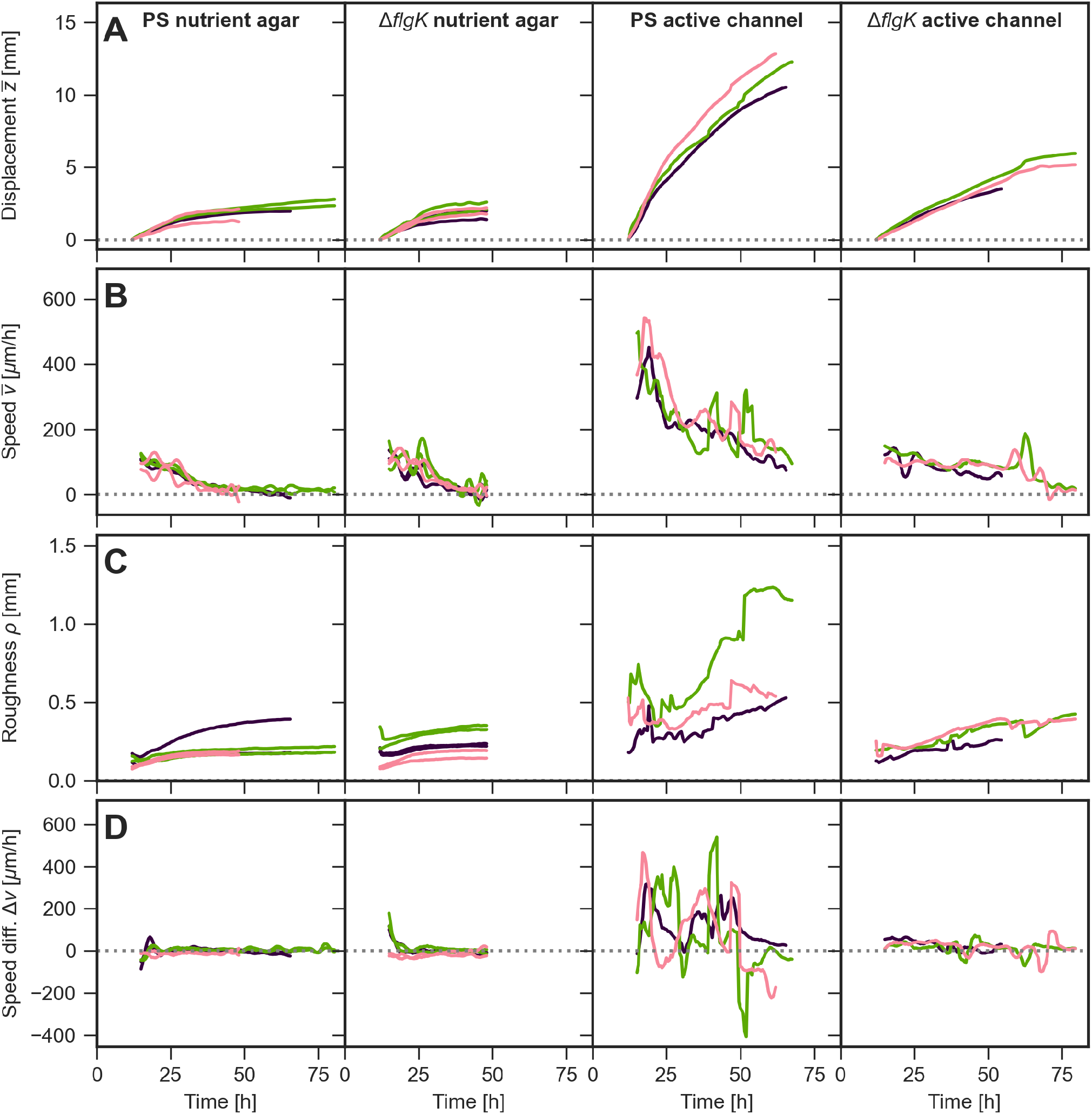
Data for individual replicates averaged in Fig. S4 on characterisation of movement of *P. aeruginosa* population fronts. A) Distance of the bacterial front from the initial position as averaged across the front over time for *P. aeruginosa* strains on nutrient agar (PS left, Δ*flgK* second from left) and on the device towards an active channel (PS second from right, Δ*flgK* right). Individual replicate experiments are indicated in purple, green and pink respectively, with two fronts for each experiment on a standard agar plate. B) Like panel A, but for mean speed of the bacterial front over time, calculated as a moving average of 6 time points over time. C) Like panel A, but for roughness of the front over time, calculated using the standard deviation of the front from its smooth contour. D) Like panel A, but for difference in front speed at the device boundaries associated with inlets and outlets, respectively.

**Figure S6:**
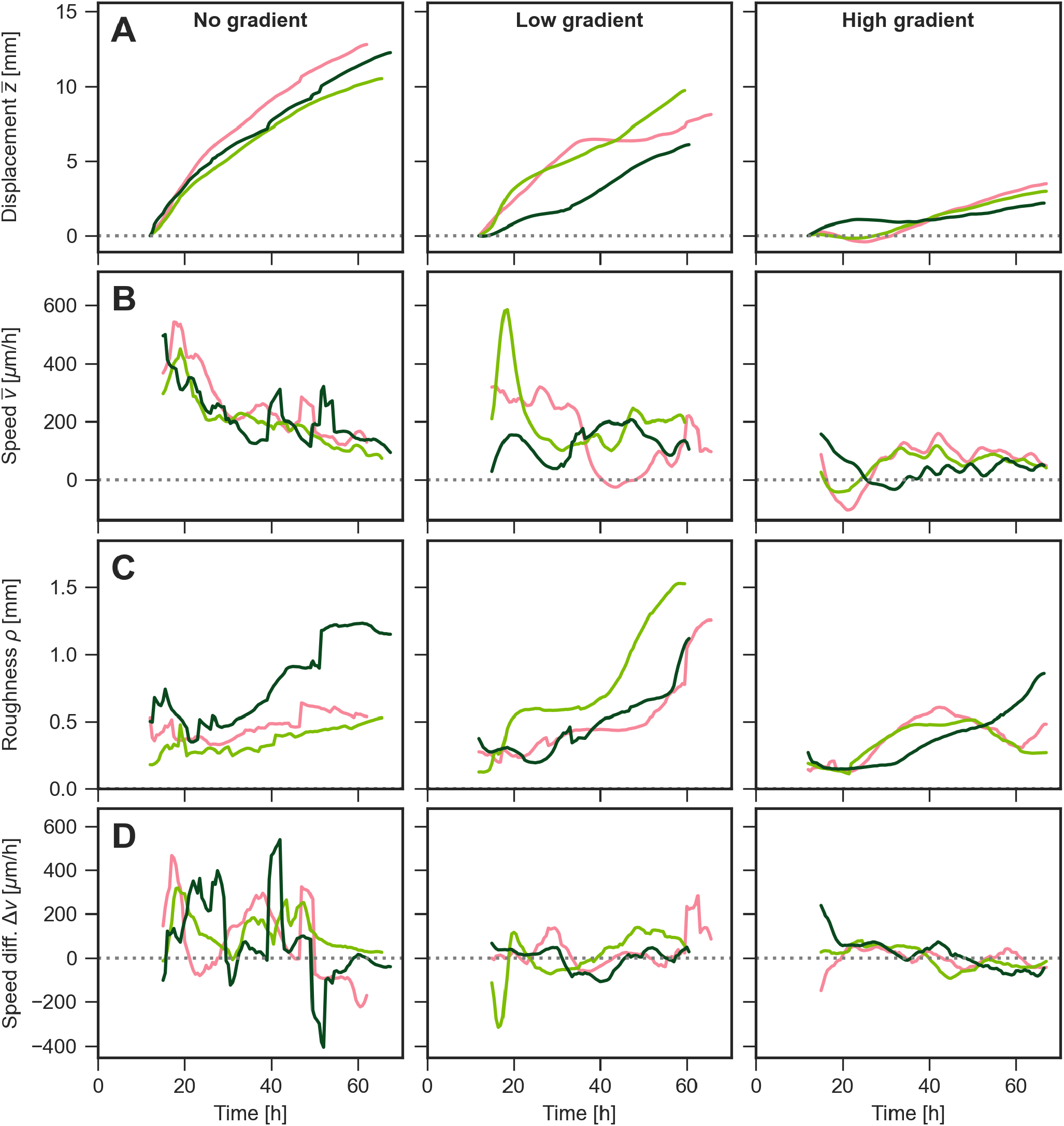
Data for individual replicates averaged in Fig. 8 on characterisation of movement of *P. aeruginosa* population fronts. A) Distance of the bacterial front from the initial position as averaged across the front over time for *P. aeruginosa* strains on nutrient agar with no antibiotic gradient (left), a shallow gradient (middle) and steep gradient (right). Individual replicate experiments are indicated in purple, green and pink, respectively. B) Like panel A, but for mean speed of the bacterial front over time, calculated as a moving average of 6 time points over time. C) Like panel A, but for roughness of the front over time, calculated using the standard deviation of the front from its smooth contour. D) Like panel A, but for difference in front speed at the device boundaries associated with inlets and outlets, respectively.

**Figure S7:**
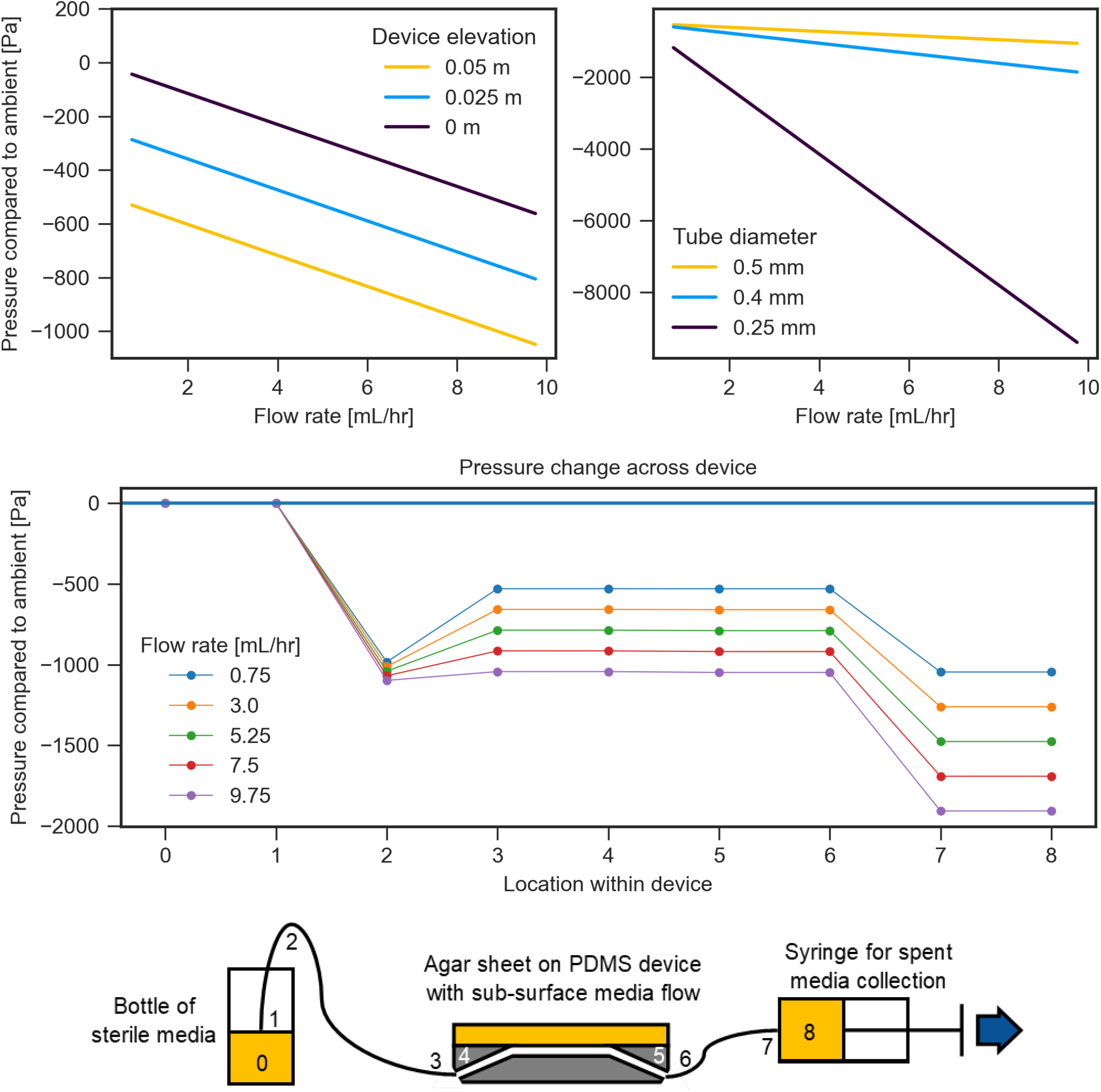
Pressure throughout the device and its periphery. (Top) Pressure within the device as a function of flow rate for different elevations of device relative to reservoir and different tube diameters. (Middle) Pressure along the device with the device elevated for different flow rates. Lines are guide to the eye only. (Bottom) Diagram of device with locations from above labelled.

**Figure S8:**
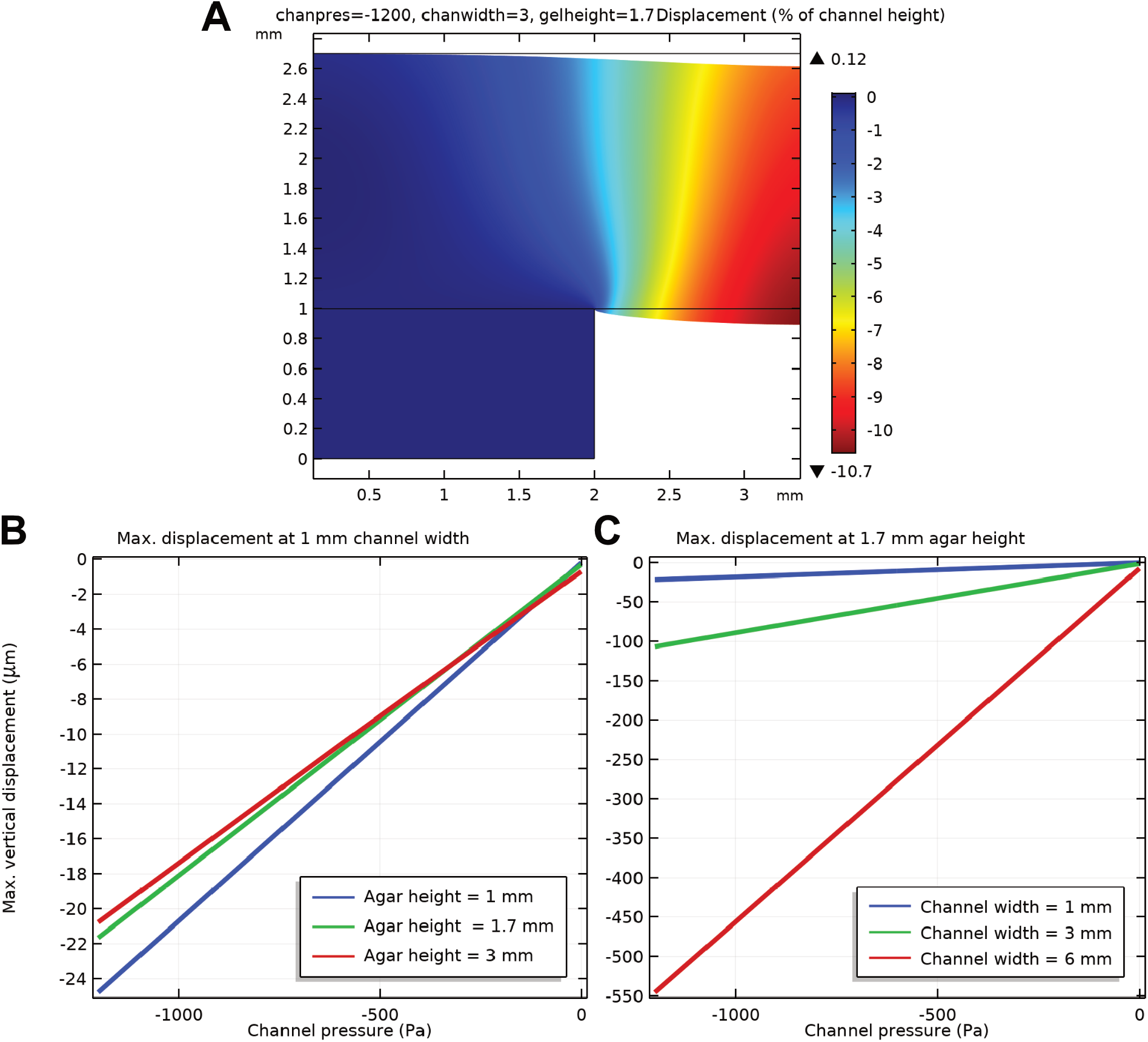
Simulated deformation of agar sheet over channel as a function of pressure in the channel. (A) Displacement over an agar sheet overhanging a channel. (B-C) Maximum displacement (at centre of channel) as a function of pressure for different agar heights and channel widths. Pressure of -1200 Pa and -600 Pa correspond to flow rates 10 mL/hr and 0.75 mL/hr as in Fig. S7.

